# GluK1 kainate receptors are necessary for functional maturation of parvalbumin interneurons regulating amygdala circuit function

**DOI:** 10.1101/2023.09.14.557707

**Authors:** Joni Haikonen, Rakenduvadhana Srinivasan, Simo Ojanen, Jun Kyu Rhee, Maria Ryazantseva, Gabija Zumaraite, Sari E. Lauri

## Abstract

Parvalbumin expressing interneurons (PV INs) are key players in the local inhibitory circuits and their developmental maturation coincides with the onset of adult-type network dynamics in the brain. Glutamatergic signaling regulates emergence of the unique PV IN phenotype, yet the receptor mechanisms involved are not fully understood. Here we show that GluK1 subunit containing kainate receptors (KARs) are necessary for development and maintenance of the neurochemical and functional properties of PV INs in the basolateral amygdala (BLA). Ablation of GluK1 expression specifically from PV INs resulted in low parvalbumin expression and loss of characteristic high firing rate throughout development. In addition, we observed reduced spontaneous excitatory synaptic activity at adult GluK1 lacking PV INs. Intriguingly, inactivation of GluK1 expression in adult PV INs was sufficient to abolish the PV phenotype, suggesting a role for GluK1 in dynamic regulation of PV IN maturation state. The PV IN dysfunction in the absence of GluK1 perturbed feedforward inhibition and long-term potentiation (LTP) in the BLA and resulted in developmentally originating changes in the glutamatergic connectivity to BLA principal neurons. Behaviorally, the absence of GluK1 from PV INs associated with hyperactivity and increased fear of novelty. These results indicate a critical role for GluK1 KARs in regulation of PV IN function across development and suggest GluK1 as a potential therapeutic target for pathologies involving PV IN malfunction.

## Introduction

Kainate receptors (KARs) are glutamate receptors with both ionotropic and metabotropic modes of action, composed of tetrameric arrangements of GluK1-5 subunits. They are widely expressed in both glutamatergic and GABAergic neurons, in pre- and postsynaptic terminals, to modulate synaptic transmission and plasticity ^1–3^. In addition, KARs are highly expressed during early postnatal development and significantly contribute to circuit development ^4–7^. Accordingly, KARs malfunction has been implicated in various neurodevelopmental and psychiatric disorders, as well as in epilepsy^8–10^. Although most of the physiological functions described for KARs in the developing circuitry are related to glutamatergic transmission, GluK1 subunit containing KARs, in particular, are also implicated in development and maturation of GABAergic interneurons and GABAergic synaptic transmission^11–14^; however, their contribution to differentiation of various interneuron subtypes remains unknown.

Parvalbumin expressing interneurons (PV INs) represent a prominent subtype of GABAergic interneurons in the brain with unique physiological properties that enable them to control local network dynamics ^15^. These interneurons reach their full electrophysiological and morphological maturity relatively late in development which, along with acquisition of extracellular components, such as perineuronal nets (PNNs), closes off the early critical period of plasticity^16,17^. Glutamatergic synaptic input is essential for activity-dependent maturation of interneurons ^18–20^. Of the various glutamate receptor subtypes, NMDA receptor mediated signaling has been shown to profoundly influence development and adult functions of the PV INs ^21–23^. PV INs also contain KARs ^24,25^ yet their roles in regulating PV IN specific functions are not well characterized.

We have recently shown that GluK1 subunit containing KARs regulate the excitability of PV INs and consequently, the function of the local disinhibitory circuit in the basolateral amygdala (BLA)^25^. Furthermore, early life stress (ELS), involving alterations in neurochemical maturation of PV INs^26^, associates with downregulation of GluK1 expression in BLA PV INs^25^. Altogether, these data suggest that KARs are in a crucial position to orchestrate interplay between glutamatergic and GABAergic systems and to drive activity-dependent maturation of PV INs.

In the present work, we used genetic tools to eliminate GluK1 expression selectively in PV INs, to study the role of GluK1 KARs in PV INs development and function, as well as its broader significance on the BLA circuitry and behavior. We show that GluK1 expression is essential for the maturation and maintenance of the PV IN phenotype. Loss of GluK1 in the PN INs associates with robust reduction in PV expression and PV IN excitability and affects feedforward inhibition and synaptic plasticity in the BLA. The amygdala circuit dysfunction observed in the mice where GluK1 was selectively ablated in the PV INs associated with behavioral changes, including hyperactivity and fear of novelty.

Taken together, these results provide vital new evidence for the critical role of GluK1 KARs in the development and maturation of PV INs, with potential implications to the variety of neurodevelopmental disorders involving perturbed PV IN function.

## Materials and methods

### Animal models

Experiments were performed using males and females between postnatal day (P) 16-70 of the following mouse lines: a conditional (floxed) line for GluK1 (*Grik1^tm^*^1c^*^/tm^*^1c^; Ryazantseva et al., 2020) and PV-Cre (JAX: 008069). PV-Cre mice were crossed with Ai14 (JAX 007914) to produce PV-TdTomato reporters and with *Grik1^tm^*^1c^*^/tm^*^1c^ to produce mice lacking *Grik1* selectively in PV interneurons (*Grik1^tm^*^1d^*^/tm^*^1d^; from here on *PV-Grik1^−/−^*). Mice from homozygote breedings were used for all experiments involving electrophysiology in juvenile age groups. For behavioral tests, functional ultrasound (fUS) imaging and some of the immunohistological stainings in adults, littermate control (*Grik1^tm^*^1c^) and *PV-Grik1^−/−^* mice that were heterozygous for Cre were used. All mice were of C57BL/6J background.

### Stereotaxic surgery and viral vectors

AAV serotype 8 viral vectors encoding for floxed EGFP under the synapsin promoter (pAAV-hSyn-DIO-EGFP (#50457)) were purchased from Addgene. rAAV-PV-EGFP-bGH polyA and rAAV-PV-CRE-EGFP-bGH polyA encoding for EGFP under the PV promoter were purchased from BrainVTA. Following craniotomical surgery, AAVs were stereotaxically injected into the BLA of adult mice under anesthesia as previously described^25^. For experiments with the juvenile cohorts, viral injections were done on neonates (P4-6) using a non-invasive method (without surgical incisions of the scalp) with minimal anesthesia, to ensure survival of pups. Viral particles were directly injected through the scalp and cranium using a stainless-steel needle (33 ga. beveled NanoFil needle, WPI). Electrophysiological or histological experiments were performed 10-14 days after viral injections on the neonatal mice, and 2-3 weeks after injections on adult mice.

### Slice preparation for electrophysiological recordings

2–3-month-old mice were anesthetized with isoflurane and quickly decapitated. The brain was extracted and immediately placed in carbogenated (95% O_2_/5%CO_2_) ice-cold *N*-Methyl-D-glucamine (NMDG) based protective cutting solution (pH 7.3-7.4) containing (in mM): 92 NMDG, 3 Na-pyruvate, 5 Na L-ascorbate, 2 thiourea, 20 HEPES, 30 NaHCO_3_, 25 glucose 1.25 NaH_2_PO_4_, 2.5 KCl, 0.5 CaCl_2_ and 10 MgSO_4_ · 7H_2_O (300-310 mOsm). For recording of virally transduced interneurons the mice were transcardially perfused with ice-cold NMDG solution prior to decapitation. The brain was glued to a stage and transferred to a vibratome (Leica VT 1200S) to obtain 300 µm thick brain slices. Slices containing the amygdala recovered for 8-12 min in the NMDG protective solution (34°C), after which they were stored in the HEPES-based solution containing (in mM): 92 NaCl, 3 Na-pyruvate, 5 Na L-ascorbate, 2 thiourea, 20 HEPES, 30 NaHCO_3_, 25 glucose, 1.25 NaH_2_PO_4_, 2.5 KCl, 2 CaCl_2_ and 2 MgSO_4_ · 7H_2_O (300-310 mOsm) at room temperature.

Juvenile mice (P16-21) were anaesthetized using isoflurane. Following rapid decapitation and opening of the cranium, the brain was placed immediately into ice-cold sucrose-based sectioning solution of the following composition (in mM): 87 NaCl, 2.5 KCl, 0.5 CaCl_2_, 25 NaHCO_3_, 1.25 NaH2PO4, 7 MgSO_4_, 75 sucrose and 25 D-glucose (300-310 mOsm). 250 µM thick coronal slices were cut in the same manner as adults and the slices recovered for 1h in ACSF containing (in mM): 124 NaCl, 3 KCl, 1.25 NaH_2_PO_4_, 26 NaHCO_3_, 15 glucose, 1 MgSO_4_. 7H_2_O, 2 CaCl_2_ (∼ 300-310 mOsm) at 34°C.

### Electrophysiological recordings

After 1–6h of recovery, the slices were placed in a submerged heated (32–34°C) recording chamber and perfused with standard carbogenated ACSF containing (in mM): 124 NaCl, 3 KCl, 1.25 NaH_2_PO_4_, 26 NaHCO_3_, 15 glucose, 1 MgSO_4_. 7H_2_O, 2 CaCl_2_ (300-310 mOsm) at the speed of 1–2ml/min. Whole-cell patch-clamp recordings were done from BLA neurons under visual guidance using patch electrodes with resistance of 2–7 MΩ. Multiclamp 700B amplifier (Molecular Devices), Digidata 1322 (Molecular Devices) or NI USB-6341 A/D board (National Instruments) and WinLTP version 2.20^27^ or pClamp 11.0 software (Molecular Devices) were used for data collection, with low pass filter (10 kHz) and sampling rate of 20 kHz. In all voltage-clamp recordings, uncompensated series resistance (Rs) was monitored by measuring the peak amplitude of the fast whole-cell capacitive current in response to 5 mV step. Only experiments where Rs <30 MΩ, and with <20% change in Rs during the experiment, were included in analysis.

### Current clamp recordings of intrinsic excitability

Whole cell recordings to characterize intrinsic excitability were done with an intracellular solution of the following composition (in mM): 135 K-gluconate, 10 HEPES, 5 EGTA, 2 KCl, 2 Ca(OH)_2_, 4 Mg-ATP, 0.5 Na-GTP, (280 mOsm, pH 7.2). Current steps of 500 ms duration and of varying amplitude (−100 to 400 pA with 25 pA increments) were injected to probe firing frequency, along with passive-membrane properties. All cells were held at resting membrane potential. Action potential (AP) half-width and amplitude of the fast afterhyperpolarizing potential (AHP) were analyzed from the 3rd spike in the train using Clampfit software (Molecular Devices).

*Spontaneous excitatory postsynaptic currents (sEPSC,)* were recorded from PV INs with the K-gluconate based pipette filling solution, following completion of current clamp recordings. The membrane potential was clamped at −70 mV and sEPSCs were recorded for 15 minutes.

*Spontaneous excitatory and inhibitory postsynaptic currents (sEPSC, sIPSC)* were recorded using a cesium-based pipette filling solution (pH 7.2, 280 mOsm) containing (in mM): 125.5 Cs-methanesulfonate, 10 HEPES, 5 EGTA, 8 NaCl, 5 QX314, 4 Mg-ATP, 0.3 Na-GTP. The membrane potential was first clamped at −70 mV and sEPSCs were recorded for 10 min. Afterwards, the membrane potential was slowly raised to 0 mV, the reversal potential of glutamatergic currents, in order to measure sIPSCs.

*Spontaneous miniature postsynaptic currents (mEPSC)* were recorded from BLA neurons using the cesium-based filling solution with 0.3% Biocytin, for later morphological analysis of spines. 1 µM of tetrodotoxin (TTX) was added to the perfusion solution. The membrane potential was clamped at −70 mV and mEPSCs were recorded for 20-30 min.

Analysis of the spontaneous synaptic events was done manually using Minianalysis program 6.0.3. EPSCs and IPSCs were identified as inward and outward currents, respectively, with typical kinetics, that were at least 3 times the amplitude of the baseline level of noise. 5 minutes of recording / cell was analyzed.

*Evoked glutamatergic and GABAergic (EPSC, dIPSC) responses* were recorded using the cesium-based filling solution. EPSCs and dIPSCs were evoked with a bipolar stimulation electrode (nickel-chromium wire), placed in the external capsule to activate the cortical inputs (0.05 Hz). The membrane potential was first held at −70 mV to record 5 min of EPSCs, after which, the cell was slowly depolarized to 0 mV to record 10-20 min of dIPSC. 10 µM CNQX was added at the end of the recording to confirm the disynaptic nature of the response (> 80% block of the IPSC amplitude). EPSC/dIPSC amplitude ratio was calculated from average of 15 responses/ condition using the WinLTP software. For experiments probing the effect of ACET on EPSCs and paired pulse (50 ms interval) ratios, a 15-minute baseline was recorded, following which 200 nM ACET was applied for 10 minutes with a consequent washout. 2-exponential fit for the EPSC decay was done using Clampfit software to separate the fast and slow components of the synaptic response. Pharmacologically isolated KAR-mediated EPSCs were recorded in the presence of 100 µM picrotoxin, 1 µM CGP55845 and 50 µM D-APV, to block GABA_A_, GABA_B_ and NMDA receptors, respectively. 50 µM GYKI53655 and 200 nM were subsequently added to antagonize AMPA and GluK1 KARs. In the presence of GYKI53655, 10 pulse / 100Hz stimulation was used to elicit a clear KAR mediated EPSC.

*Field recordings* were done using an interface chamber (32–34°C). fEPSPs were evoked with a bipolar stimulation electrode (nickel-chromium wire) placed in the external capsule, while the recording electrode (resistance 1–3 MΩ, filled with standard ACSF) was placed in the lateral amygdala (LA). The stimulation intensity was adjusted to produce a half-maximal response size. After a stable baseline, LTP was induced using theta-burst stimulation (TBS) (6 bursts of 8/100 Hz at 5 Hz), repeated 3 times with 5 min intervals. After TBS, synaptic responses were recorded for 60 min. Analysis was done using the WinLTP software, averaging 3 responses per timepoint. Averages to evaluate the level of synaptic potentiation were calculated over a stable 10 min period, just before TBS and at the end of the recording.

### Immunohistochemistry

For parvalbumin (PV) immunostaining, P21 and P60 mice were intracardially perfused with PBS and 4% PFA under deep anesthesia. The brains were extracted and placed in 4% PFA overnight at 4°C, after which they were washed twice with PBS. 40 µM thick coronal sections were cut using a vibratome (Leica VT 1200S) and placed in 12-well well plates. The sections were incubated in a blocking solution (10% goat serum, 1% BSA and 0.5% Triton-X in PBS), in room temperature (RT), for 1h, then washed with a wash buffer (1% goat serum, 1% BSA and 0.5% Triton-X in PBS) for 20 min in RT, and finally in primary antibodies diluted into the wash buffer (1:1000 PV 235 mouse (SWANT)) overnight in 4°C. The next day, the sections were washed with the wash buffer 2 x 20 min in RT, incubated in secondary antibodies diluted into the wash buffer (1:1000 Alexa 488 goat anti-mouse (Life Technologies)) for 1h in RT, and finally washed with PBS 3 x 20 min in RT, and mounted on to microscope slides using Vectashield mounting medium. The stained sections were imaged with Zeiss Axio Imager.M2 microscope equipped with ApoTome 2 structured illumination slider, using PlanApo ×10 objective, CMOS camera (Hamamatsu ORCA Flash 4.0 V2) and Zen Blue software. All sections were imaged with the same exposure time for integrated optical density (IOD) analysis which was calculated using ImageJ software^28^.

*Spine density analysis* was done *post hoc* for biocytin filled amygdala neurons. Sections were fixed overnight in 4% PFA in 4°C, washed with PBS and placed in a permeabilization solution (0.3% Triton-X in PBS) overnight in 4°C. 1:500 streptavidin Alexa 568 (Life Technologies) was added (overnight in 4°C), after which the sections were mounted on microscope slides using ProLong™ Gold Antifade Mountant (Thermofisher). Dendritic spines were imaged using a LSM Zeiss 710 confocal microscope (alpha Plan146 Apochromat 63x/1.46 OilKorr M27 objectives) with resolution of 15.17 pixels/µm and Z-stack interval of 0.5 µm. Spine density was calculated from the first branch (length 60-100 µm) of the dendritic tree. Analysis was done manually using ImageJ software.

### Functional ultrasound imaging

Resting state functional connectivity between medial prefrontal cortex (mPFC) and basolateral amygdala (BLA) in P21 and P60 littermate control and PV-*Grik1^−/−^* mice was measured using Iconeus One functional ultrasound imaging system. The animals were anesthetized using medetomidine (1 mg/kg) + ketamine (75 mg/kg), after which the head fur was shaven off. The animal head was fixed in a stereotaxic frame with heat pad (37°C) and ultrasound gel was spread on the scalp. The probe was controlled using the IcoScan (v. 1.3.1) software, first placing it near the scalp and then locating the posterior cerebral arteries (PCA). A coronal mapping (Angio3D) scan was performed from the PCA to the mPFC with a slice interval of 0.1 mm. The scan was uploaded to IcoStudio (v. 1.2.1) software and was mapped using the automated mapping system in the software. Regions of interest were selected (mPFC, BLA, hippocampus) and two markers were placed on the mapping, one in mPFC and the other in BLA. Probe coordinates were computed based on these markers, to align all areas of interest in a single plane. The coordinates were manually uploaded to the IcoScan software, and a 10 min 2D functional ultrasound scan was performed. After the scan, the animals from the P21 cohort were placed into a heat chamber (37°C) and injected with atipamezole (0.5 mg/kg) for recovery for later P60 measurements. IcoStudio software was used to compute a functional connectivity matrix, based on the functional ultrasound scan. Baseline correction and a 0.2Hz low pass filter were applied to the final results. For analysis, the matrix values (Pearson’s correlation coefficients) were Fisher transformed.

### Behavioral tests

#### Elevated plus-maze (EPM)

The maze consisted of two open arms (30 x 5 cm) and two enclosed arms (30 x 5 cm) connected by central platform (5 x 5 cm) and elevated to 40 cm above the floor. Open arms were surrounded by 0.5 cm high edges to prevent accidental falling of mice. The floor of each arm was light grey and the closed arms had transparent (15 cm high) side- and end-walls. The illumination level in all arms was ∼150 lx. The animal was placed in the center of the maze facing one of the closed arms and observed for 5 minutes. The trials were recorded with Ethovision XT13 videotracking software (Noldus Information Technology) to measure the time spent in different zones of the maze.

#### Open field (OF)

The mice were released in the corner of novel open field arena (white floor, transparent walls, 30 x 30 cm, Med Associates). Horizontal activity was recorded by infrared sensors for 30 minutes (light intensity ∼150 lx). Peripheral zone was defined as a 6 cm wide corridor along the wall.

#### Fear conditioning (FC)

The experiments were carried out with a computer-controlled fear conditioning system (TSE Systems). Training was performed in a transparent acrylic cage (23 × 23 × 35 cm) within a constantly illuminated (∼ 100 lx) fear conditioning box. A loudspeaker provided a constant, white background noise (68 dB) for 120 s followed by 10 kHz tone (CS, 76 dB, pulsed 5 Hz) for 30 s. During conditioning, the tone was terminated by a footshock (US, 0.6 mA, 2 s, constant current) delivered through a stainless steel grid-floor (bar Ø 4 mm, distance between the bars 10 mm). Two CS-US pairings were separated by a 30 s pause. Contextual memory was tested 24 h after the training. The animals were returned to the conditioning box and total time of freezing (defined as an absence of any movements for more than 2 s) was measured through infrared light beams scanned continuously with a frequency of 10 Hz. The CS was not used during this time. Memory for the CS (tone) was tested 2 h later in a novel context. The new context was a similarly sized acrylic box with black non-transparent walls and smooth floor. A layer of wood chips (clean bedding material) under the floor provided a novel odour to the chamber. After 120 s of free exploration in a novel context the CS was applied for additional 120 s and freezing was measured as above.

#### Statistical analysis

All statistical analyses were done on raw (not normalized) data using Sigma Plot or Prism GraphPad software. All data were first assessed for normality and homogeneity of variance and the statistical test was chosen accordingly. Differences between two groups were analyzed using two-tailed t-test, paired t-test, Wilcoxon test, Mann–Whitney rank-sum test, one-way repeated measures ANOVA, or mixed effects model (REML), as appropriate. Two-way ANOVA with Holm–Sidak post-hoc comparison and Kruskal Wallis test was used to compare effects of sex and genotype or age and genotype. All the pooled data are given as mean ± SEM.

## Results

### GluK1 subunit containing KARs contribute to glutamatergic synaptic transmission to PV interneurons in the juvenile BLA

Our previous research has shown the presence of functional GluK1 subunit containing KARs in PV INs in the adult mouse BLA^25^. To confirm the presence of GluK1 KARs in PV INs also in the developing amygdala, we recorded excitatory postsynaptic currents (EPSCs) from PV INs of 16-18 day old PV-tdTomato mice in response to stimulation of cortical afferents, and investigated their sensitivity to ACET, an antagonist selective for GluK1 subunit containing receptors^29^. Application of ACET (200 nM) resulted in significant reduction in the EPSC peak amplitude (p = 0.0002, paired t-test), but had no effect on paired-pulse ratio (Figure 1A, B). These data indicate that GluK1 containing KARs regulate glutamatergic drive onto PV INs. Furthermore, the lack of effect of ACET on paired-pulse ratio suggest that these KARs are located postsynaptically and contribute to the EPSC.

**Figure 1.**
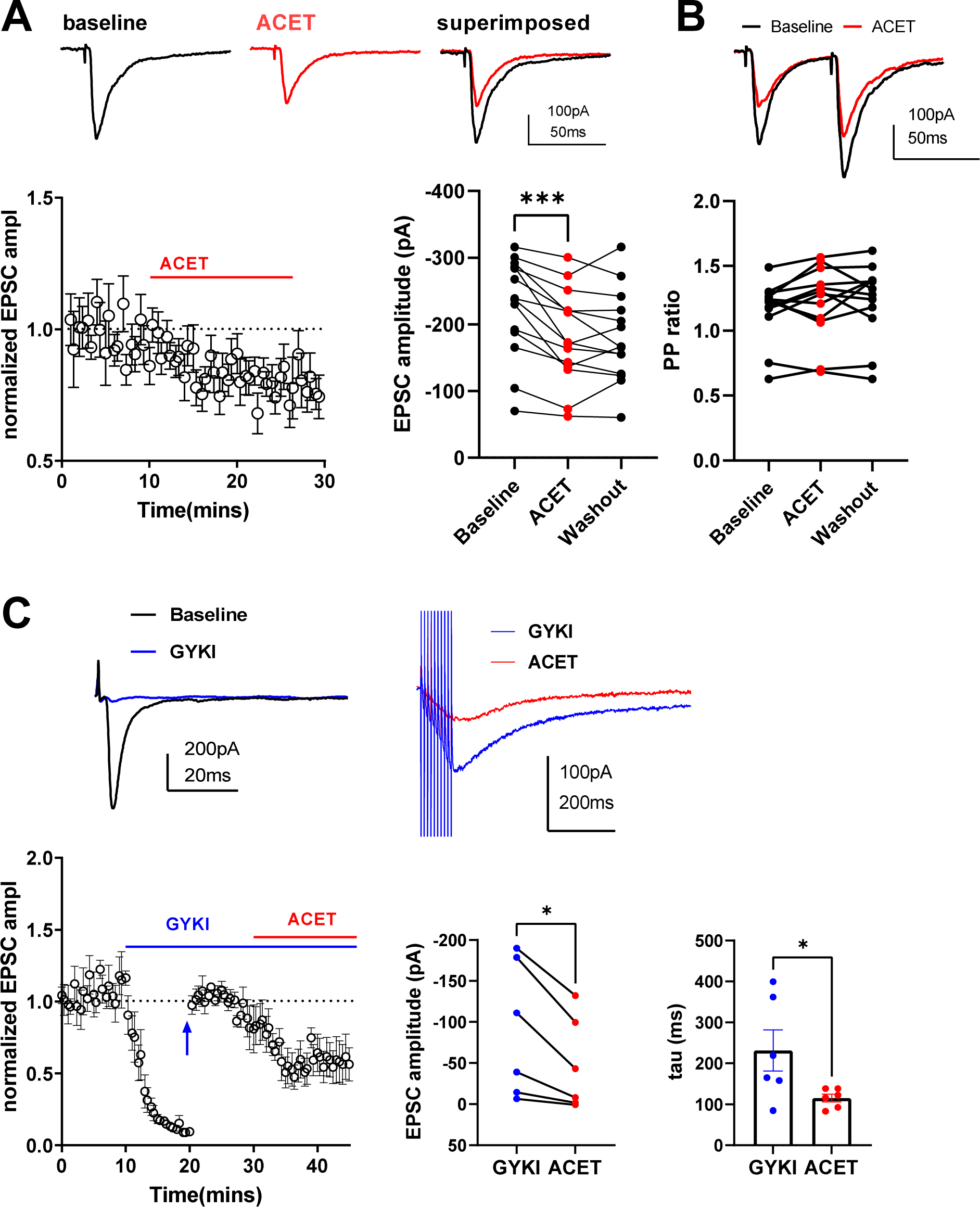
GluK1 subunit containing KARs contribute to glutamatergic synaptic transmission at the cortical inputs to PV INs in the juvenile BLA. **A.** Example traces and quantified data illustrating the effect of 200 nM ACET on the EPSC amplitude, in BLA PV INs of 16-18 day old PV-tdTomato mice (n = 14 cells, 4 animals). ***p < 0.001, paired t-test. **B.** Paired pulse (PP) ratio of EPSCs before and after ACET application, from the same cells as in A. **C.** Example traces of pharmacologically isolated EPSCs and time course plot of normalized EPSC amplitude (n = 6 cells, 3 animals). Baseline AMPA-KAR mediated EPSCs were recorded in the presence of 100 µM picrotoxin, 1 µM CGP55845 and 50 µM D-APV, to block GABA_A_, GABA_B_ and NMDA receptors, respectively, with subsequent application of 50 µM GYKI53655, a selective antagonist of AMPA receptors. 10 min after wash-in of GYKI53655 (Blue arrow), stimulation intensity was increased, 10 pulse (100Hz) stimulation was commenced to visualize KAR mediated component of the EPSC, and the EPSC amplitudes were normalized to the new baseline. Application of 200 nM ACET resulted in significant inhibition of the pharmacologically isolated KAR-EPSC. *p= 0.0176, paired t-test and * p= 0.0312, Wilcoxon test for amplitudes and decay time constant comparisons, respectively.

To validate the postsynaptic localization of the GluK1 KARs, we went on to pharmacologically isolate the KAR-mediated component of the EPSC in the juvenile BLA PV INs. Application of GYKI53655 (50µM) fully blocked the fast component of the AMPA-KAR EPSC, recorded in the presence of GABA_A_, GABA_B_ and NMDA receptor antagonists. Increasing the stimulation intensity in the presence of GYKI53655 revealed a low-amplitude, slowly decaying KAR mediated EPSC that was significantly reduced in response to application of ACET (200 nM) (p= 0.0112, paired t-test)(Figure 1C).

### GluK1 regulates parvalbumin expression and the density of PV interneurons in the BLA

Having confirmed that GluK1 KARs contribute to synaptic transmission onto PV INs in the BLA already at juvenile stage, we investigated how the absence of GluK1 in PV INs affects their maturation via immunohistochemistry at two time points, P21 and P60. To this end, we used conditional knockout mice lacking GluK1 KAR expression selectively in PV INs (*PV*-*Grik1^−/−^*) and analyzed PV+ cell density and the intensity PV labelling, commonly used as a measure of PV IN neurochemical maturation (Figure 2A). Comparison of PV+ cell density in the BLA between control (*Grik1^tm^*^1c^) and *PV*-*Grik1^−/−^* male mice revealed a significant effect of genotype (F_(1, 11)_ = 46.951, p < 0.001, 2-way ANOVA). Post-hoc test indicated that PV+ cell density was lower in both adult and juvenile *PV*-*Grik1^−/−^* as compared to controls (Figure 2A). Analysis of the integrated optical density (IOD) of PV labeling from the same samples demonstrated a significant effect of age (F_(1, 11)_ = 31.985, p < 0.001, 2-way ANOVA) and genotype (F_(1, 11)_ = 129.053, p < 0.001, 2-way ANOVA). Post-hoc comparisons revealed that PV+ cell IOD was significantly lower in both adult and juvenile *PV*-*Grik1^−/−^* mice, as compared to their respective controls (Figure 2A).

**Figure 2.**
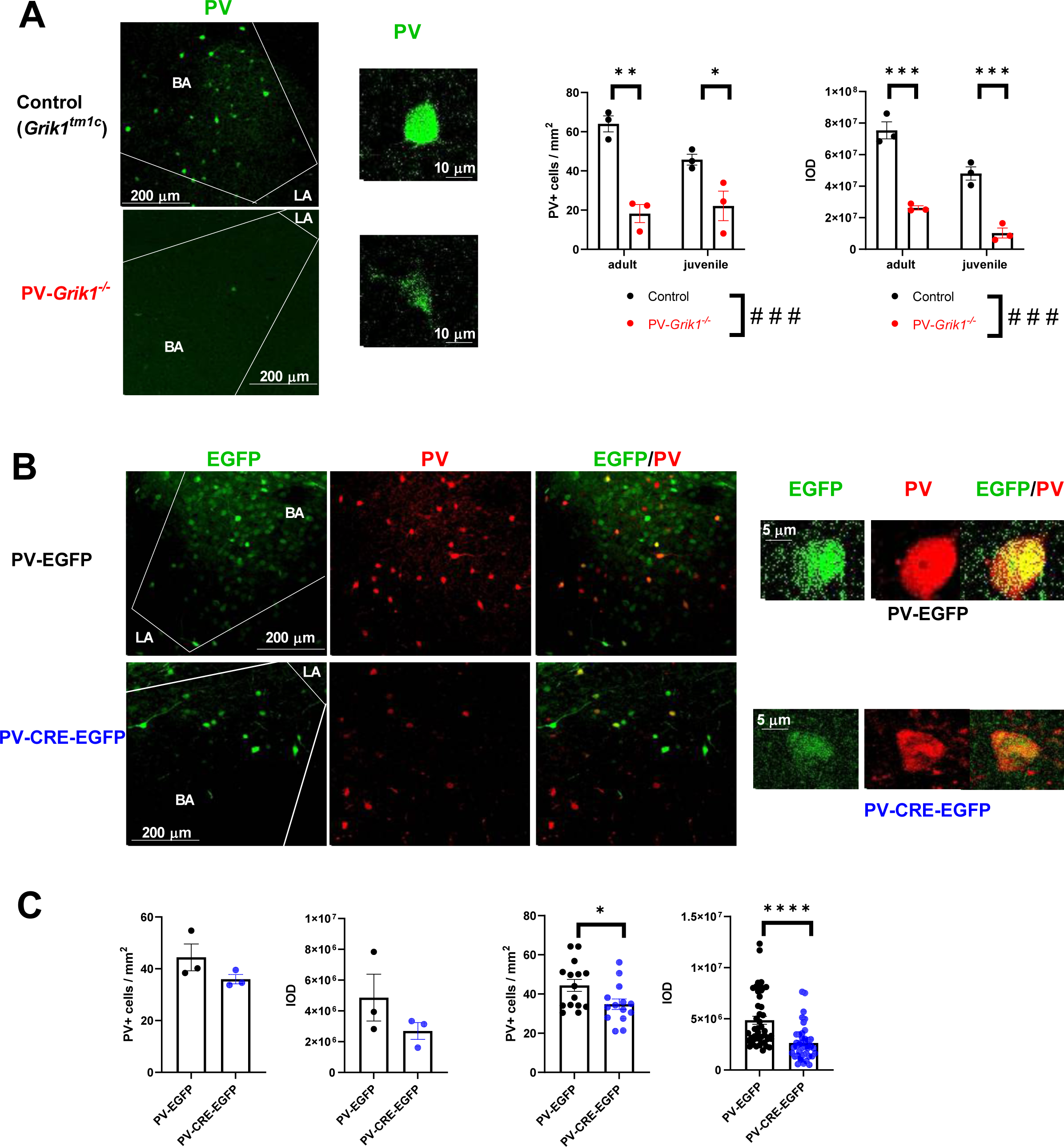
GluK1 lacking PV interneurons are characterized by low PV expression. **A.** Example images of PV (green) immunostaining in the BLA of adult control (top row) and *PV-Grik1^−/−^* (bottom row) mice and enlarged inset of a single control (top) and *PV-Grik1^−/−^* (bottom) PV+ cell. On the right, quantification of the PV+ cell density and integrated optical density (IOD) of the PV labeling, in both adult and juvenile BLA. The data consists of 3 animals per group and 5 - 8 slices per animal. *p < 0.05; ***p < 0.001; Holm-Sidak, ### p < 0.001; 2-way ANOVA. **B.** Example pictures of PV (red) immunostainings in the BLA of adult *Grik1^tm^*^1c^ mice, transduced with AAV viruses encoding for PV-EGFP (green; top row; control), or PV-CRE-EGFP (green, bottom row). On the right, enlarged example pictures of single EGFP expressing PV+ cells in both groups. **C.** Quantification of PV+ cell density and IOD from PV-EGFP and PV-CRE-EGFP injected adult *Grik1^tm^*^1c^ mice. The data are collected from 3 animals per group and 5-8 slices per animal, and are shown for both, average per animal (left) and average per slice. *p < 0.05; ***p < 0.001; one-way repeated measures ANOVA.

We also investigated how GluK1 ablation affected PV expression and density in females vs males. These experiments were done only in the adult age group, using littermate control and *PV*-*Grik1^−/−^* mice that were heterozygous for *Cre*. Interestingly, unlike the *Cre* homozygotes used in the first stainings (Figure 2A), the PV+ cell density and labelling intensity were only moderately affected in *PV*-*Grik1^−/−^* mice that were heterozygous to *Cre* (Supplementary Figure 1), indicating that there seems to be a dose dependent effect of CRE on how strongly PV INs are affected by the absence of GluK1. However, the genotype effect between *PV*-*Grik1^−/−^* and control mice was not significantly different between males and females, for either the PV+ cell density (males, 81 ± 9 %, females 84 ± 12 % of control), or PV labeling intensity (males 76 ± 7 %, females 79 ± 6 % of control; Supplementary Figure 1).

Next, we wanted to see whether a local inactivation of GluK1 expression in PV INs in adulthood is sufficient to produce the perturbed phenotype. We injected AAV viral vectors encoding for PV-EGFP (control) or PV-CRE-EGFP (*PV*-*Grik1* KO) to the BLA of a conditional (floxed) mouse line for GluK1 (*Grik1^tm^*^1c^) and used them for PV immunostainings 2-3 weeks after the injection. Since the specificity of these viral vectors for PV INs was less than ideal (EGFP+PV+ cells / EGFP+ cells, PV-EGFP: 11.81 ± 1.74%, PV-CRE-EGFP: 58.42 ± 3.25 %), only cells that were positive for both EGFP and PV were selected for IOD analysis. Male mice infected with PV-CRE-EGFP had significantly lower PV+ cell density (F_(1, 28)_ = 5.544, p = 0.026) and IOD in the EGFP-positive PV+ cells (F_(1, 86)_ = 38.845, p < 0.001) in the BLA as compared to controls (Figure 2B and C).

Taken together, these results show that GluK1 KARs profoundly affect maturation of the PV interneurons in the BLA.

### GluK1 is necessary for the high-frequency firing phenotype of PV interneurons in the BLA

Having observed the profound neurochemical alterations in our immunohistochemical experiments, we set out to investigate whether absence of GluK1 expression affected physiological functions of PV INs. To be able to identify PV INs, AAV-viruses encoding for *Cre*-dependent EGFP were injected to the BLA of *PV*-*Grik1^−/−^* and control (*PV-Cre)* mice 2-3 weeks before *in vitro* electrophysiological experiments. Current clamp recordings of EGFP positive PV INs revealed that their firing rate in response to depolarizing current steps was significantly lower in the *PV*-*Grik1^−/−^*mice in comparison to controls (p < 0.0001, mixed effects model)(Figure 3A), consistent with a loss of their characteristic high frequency firing phenotype. This effect was highly significant in both males and females (Supplementary Figure 2A), when analysed separately. Furthermore, in the adult *PV*-*Grik1^−/−^* mice, the rheobase of the PV INs was significantly higher (p = 0.035, Mann Whitney test) and input resistance was lower (p < 0.0001, Mann-Whitney test) than in controls, while resting membrane potential remained the same across both the groups and sexes (Figure 3B). Next, we investigated if inactivation of GluK1 expression in the adulthood is sufficient to produce this profound loss of PV IN excitability. Since this phenotype was seen in both males and females in the *PV*-*Grik1^−/−^* adults, we focused only on males in this dataset. Targeted ablation of GluK1 expression in the BLA was done using the PV-EGFP and PV-CRE-EGFP viral vectors as above; however, as these viral vectors were not fully specific to PV INs, we used additional criteria to exclude prospective principal neurons. First, only those EGFP+ neurons having a somatic diameter of less than 15 μm, that were spherical or fusiform in appearance, were selected for recording under visual guidance ^30,31^. Furthermore, cells with action potential halfwidth >1 ms, and spike frequency adaptation (spike interval ratio < 0.7, for 1st-2nd / 2nd-3rd AP), were excluded at the analysis stage (representing 17 % of recorded neurons)^32,33^.

**Figure 3.**
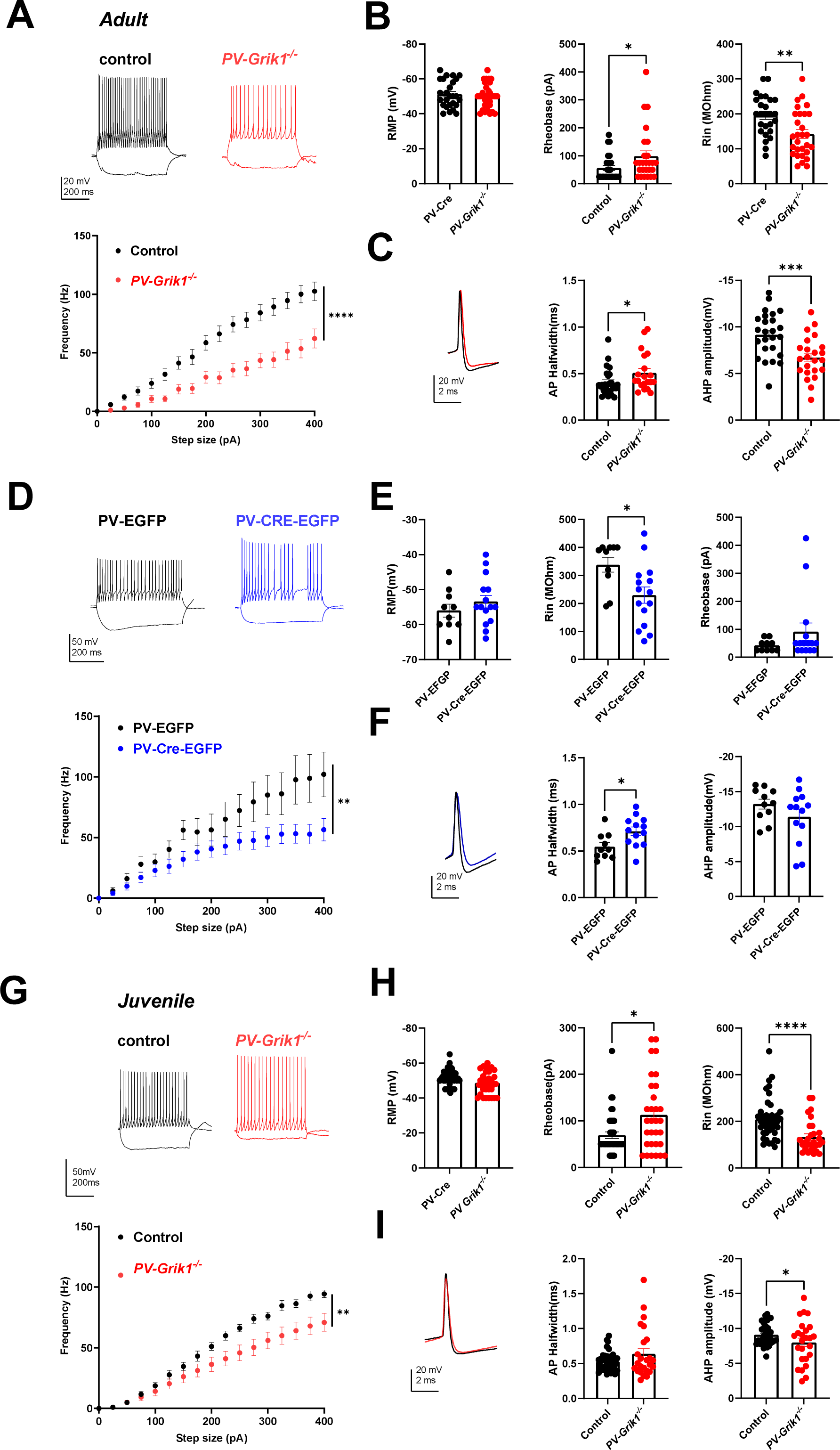
Absence of GluK1 results in low PV IN excitability across and beyond development. **A.** Example traces illustrating PV INs response to depolarizing step (200 pA) in adult control (*PV*-*Cre,* black traces) and *PV*-*Grik1^−/−^* (red traces) mice. Quantification of the firing frequency of PV INs in response to depolarizing current steps, including data from both males and females (control: n = 25 cells, 8 animals, *PV*-*Grik1^−/−^*: n = 26 cells, 7 animals) ****p < 0.0001, Mixed effects model. Data from males and females separately are shown in Supplementary Figure 2. **B.** Pooled data on resting membrane potential (RMP), rheobase and input resistance (Rin), for the same recordings as in A. *p < 0.05; **p < 0.01; ***p < 0.001, Mann-Whitney. **C.** PV interneuron action potential waveform from adult control (*PV*-*Cre,* black traces) and *PV*-*Grik1^−/−^* (red traces) mice, and pooled data for AP half-width and fast after-hyperpolarization (AHP) current amplitudes in both genotypes. Data is from the same cells as in A. **D.** Example traces and averaged data on the firing rate of PV-EGFP (black traces) and PV-CRE-EGFP (blue traces) expressing PV INs in adult male mice (control: n = 10 cells, 3 animals, PV*-Grik1* KD: n = 15 cells, 3 animals) *p < 0.009, Mixed effects model. **E.** Quantification of the membrane properties for the same cells as in D. *p < 0.05; Mann-Whitney. **F.** Action potential waveform and quatification of AP half-width and fast after-hyperpolarization (AHP) amplitudes for the same cells as in D. *p < 0.05; Mann-Whitney. **G.** Similar data as in A, from juvenile mice (control: n = 41 cells, 8 animals, *PV*-*Grik1^−/−^*: n = 30 cells, 8 animals) *p < 0.009, Mixed effects model **H, I.** Quantification of the membrane properties and example traces of the action potential waveform for the same cells as in G.. *p < 0.05; **p < 0.01; ***p < 0.001, ****p < 0.0001, Mann-Whitney.

*PV*-*Grik1* KO cells, identified using the above criteria, displayed low excitability, similar to that seen in the *PV*-*Grik1^−/−^*mice. Thus, the firing frequency in response to depolarizing current steps was significantly lower in *PV*-*Grik1* KO cells than in controls (p < 0.0092, mixed effects model, Figure 3D). *PV*-*Grik1* KO cells also had a significantly lower input resistance (p = 0.0228, Mann Whitney test, Figure 3E) than controls. Although there was a trend of higher rheobase values in the *PV*-*Grik1* KO vs control PV INs, this was not statistically significant and resting membrane potentials were unaffected (Supplementary Figure 3E).

To examine whether the loss of excitability in GluK1 lacking PV INs is already present during development, we conducted current clamp recordings of BLA PV INs from juvenile *PV*-*Grik1^−/−^* mice. Our experiments revealed that the firing frequency of PV INs was significantly lower in *PV*-*Grik1^−/−^* mice as compared to controls when all the data was pooled together (p < 0.0093 Mixed-effects model) (Figure 3G). However, the effect of GluK1 elimination on PV IN excitability was not significant in juvenile females (Supplementary Figure 2B), possibly because of a difference in developmental trajectories and timings between sexes. Moreover, we observed higher rheobase (p = 0.019, Mann Whitney) and lower input resistance (p < 0.0001, Mann Whitney), across both sexes in *PV*-*Grik1^−/−^* PV INs (Figure 3H; Supplementary Figure 2B).

For further insight on the active membrane properties of PV INs, we also compared action potential (AP) half width and the amplitude of the fast afterhyperpolarization (AHP) between the genotypes. The AP-halfwidth decreases in PV INs as a consequence of development, accompanying the augmentation of their firing rate ^34,35^. Accordingly, we observed a significant decrease in AP halfwidth in the control PV INs from juvenile to adult stage (p < 0.0001, Mann-Whitney test, Supplementary Figure 2C). In adult *PV*-*Grik1^−/−^* PV INs, but not at the juveniles, the AP-halfwidth was significantly higher as compared to controls (p = 0.0410, Mann Whitney test; Figure 3C,I), suggesting impaired maturation of PV INs. Indeed, there was no difference in AP halfwidth between juvenile and adult *PV*-*Grik1^−/−^* PV INs (p = 0.2155, Mann-Whitney test, Supplementary Figure 2A). In addition, we observed that AP-halfwidth was higher in the adult *PV*-*Grik1* KO cells as compared to controls (p = 0.0146, Mann Whitney test, Figure 3F), suggesting that GluK1 KARs play a vital role in maintaining the fast action potential kinetics of adult PV INs.

After-hyperpolarization (AHP) current is an electrophysiological signature that distinguishes PV INs from other interneurons ^15,36–38^. Our experiments revealed that the amplitude of the fast AHP current was significantly lower in both, adult and juvenile *PV*-*Grik1^−/−^* PV INs of both sexes (p = 0.0006, two tailed t test and p = 0.0332, Kolmogorov-Smirnov test, respectively) but not significantly lower in *PV*-*Grik1* KO neurons, as compared to their respective controls (Figure 3C, F, I).

These results indicate a crucial role of GluK1 in maintaining PV IN excitability and electrophysiological characteristics (such as the high frequency firing) across development, extending beyond the developmental windows of plasticity.

### GluK1 ablation perturbs glutamatergic inputs to PV interneurons

We next examined whether absence of GluK1 affected glutamatergic input to PV INs by conducting whole cell voltage clamp recordings of spontaneous excitatory postsynaptic currents (sEPSCs) in BLA PV INs, identified by expression of *Cre*-dependent EGFP in *PV*-*Grik1^−/−^* and control (*PV-Cre)* mice. These recordings revealed significantly lower frequency and amplitude of sEPSCs in the adult *PV-Grik1^−/−^* PV INs as compared to controls (p = 0.0005, Mann Whitney test and p=0.0004, two-tailed t test, respectively) (Figure 4A). Interestingly, these effects seemed to be stronger in females than in males (Supplementary Figure 3A). Similar changes in glutamatergic input to PV INs were also observed when GluK1 was locally eliminated in the adults, by using the AAV-PV-CRE-EGFP viral vectors in *Grik1^tm^*^1c^ mice. The sEPSC frequency, but not the amplitude was significantly lower in *PV*-*Grik1* KO neurons in comparison to controls (p = 0.0003, Mann Whitney) (Figure 4B). In these recordings, putative principal neurons were excluded from the analysis using the same visual and electrophysiological criteria as defined for the data in Figure 3.

**Figure 4.**
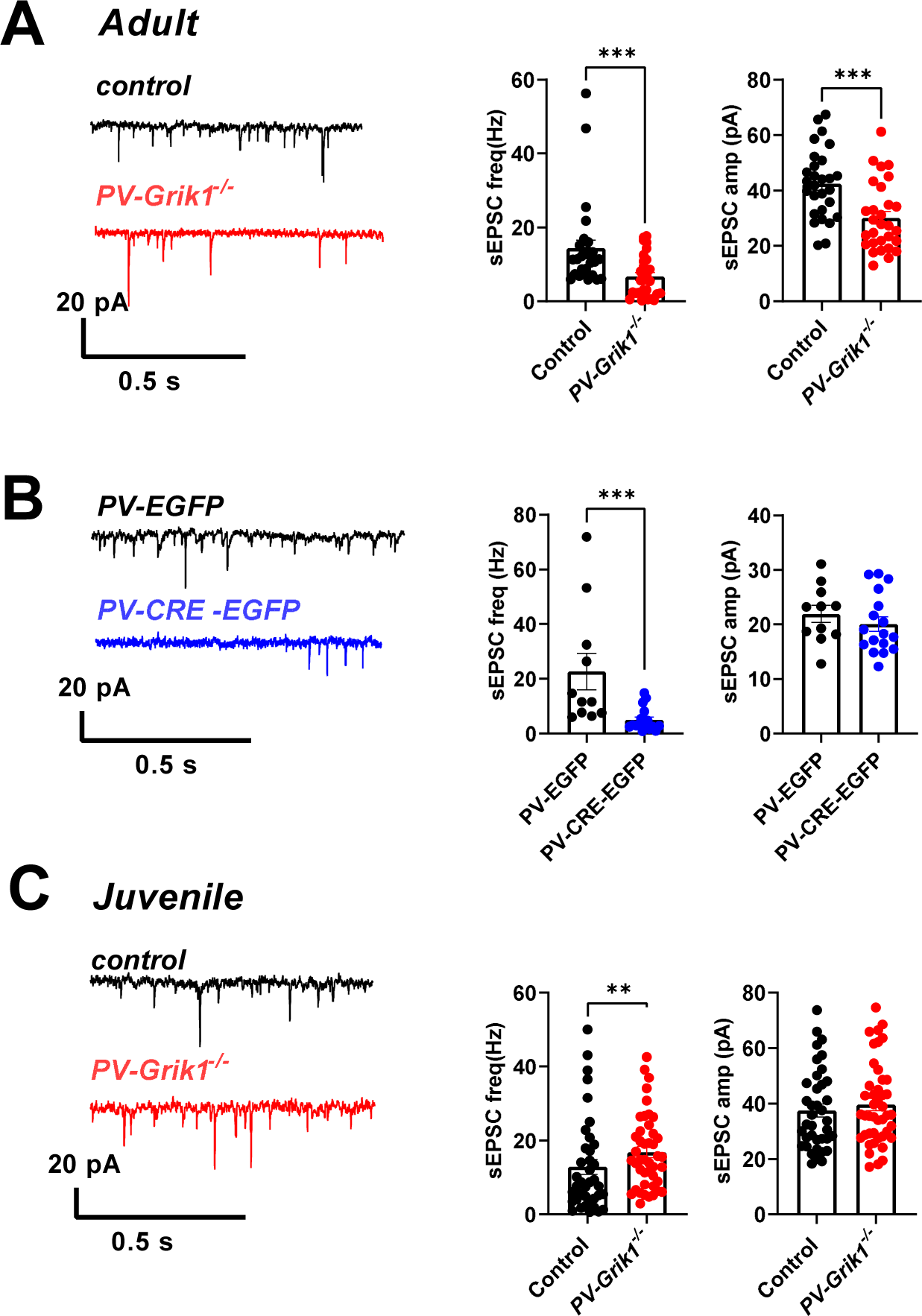
Absence of GluK1 perturbs glutamatergic input to PV INs across and beyond development. **A.** Example traces (left) and quantification (right) of PV INs sEPSC frequency (males and females pooled) and amplitude in adult control (*PV-Cre,* black traces) and *PV*-*Grik1^−/−^* (red traces) mice (control: n = 28 cells, 8 animals *PV*-*Grik1^−/−^*: n = 29 cells, 7 animals) ***p < 0.001; **** p < 0.0001, Mann-Whitney. **B.** Example traces (left) and quantification (right) of sEPSC frequency and amplitude in control (PV-EGFP, black trace) and *Grik1* knockout (PV-CRE-EGFP, blue trace) PV INs (PV-EGFP: n = 11 cells, 3 animals, PV-CRE-EGFP: n = 17 cells, 3 animals) ***p < 0.001; Mann-Whitney. **C.** Example traces (left; control: black traces, *PV*-*Grik1^−/−^:* red traces) and quantification (right) of PV INs sEPSC frequency and amplitude in juvenile (males and females pooled) control (n = 38 cells, 9 animals) and *PV*-*Grik1^−/−^* (n = 34 cells, 10 animals) mice **p < 0.01, Mann-Whitney.

Interestingly, in contrast to the adults, sEPSC frequency was significantly higher in the juvenile (P16-18) *PV-Grik1^−/−^* PV INs as compared to controls (p=0.0096; Mann Whitney test; Figure 4C), in both sexes (Supplementary Figure 3B). sEPSC amplitudes were not different between the groups.

These data indicate the GluK1 expression in the adult stage is necessary to maintain glutamatergic input to PV INs. In juveniles this effect was not observed, possibly because of efficient homeostatic compensatory mechanisms during the early plasticity window.

### Elimination of GluK1 in PV interneurons affects development of the synaptic circuitry in the BLA

Since the absence of GluK1 KARs in PV INs led to alterations in their neurochemical and functional maturation, we investigated the broader consequences of this on BLA circuitry by recording excitability and synaptic inputs in BLA principal neurons (PNs). We decided to focus our efforts solely on males, as the changes in PV IN excitability, AP kinetics and membrane properties in *PV*-*Grik1^−/−^* mice were similar in both sexes. Whole cell recordings indicated that the firing frequency of the adult BLA PNs in response to depolarizing steps was not different between the control and *PV*-*Grik1^−/−^*groups (F_(1, 131)_ = 0.9559, p = 0.33, mixed effects model; Figure 5A). Also, we detected no differences between the genotypes in the ongoing GABAergic inhibition in either age group, when assessed by spontaneous inhibitory postsynaptic currents (sIPSC) (F_(1, 46)_ = 0.236, p = 0.63, 2-way ANOVA) (Figure 5B). Interestingly, there was a significant genotype effect on the frequency of sEPSCs, reflecting the ongoing glutamatergic input to BLA PNs (F_(1, 45)_ = 14.886, p < 0.001, 2-way ANOVA). Post-hoc comparisons revealed that sEPSC frequency in adult *PV*-*Grik1^−/−^*mice was significantly lower as compared to controls (Figure 5B). A similar trend was visible in juvenile animals, yet this effect was not statistically significant (Figure 5B). No differences between the genotypes were detected in sEPSC or sIPSC amplitudes, either in adult or juvenile mice (Supplementary Figure 3C).

**Figure 5.**
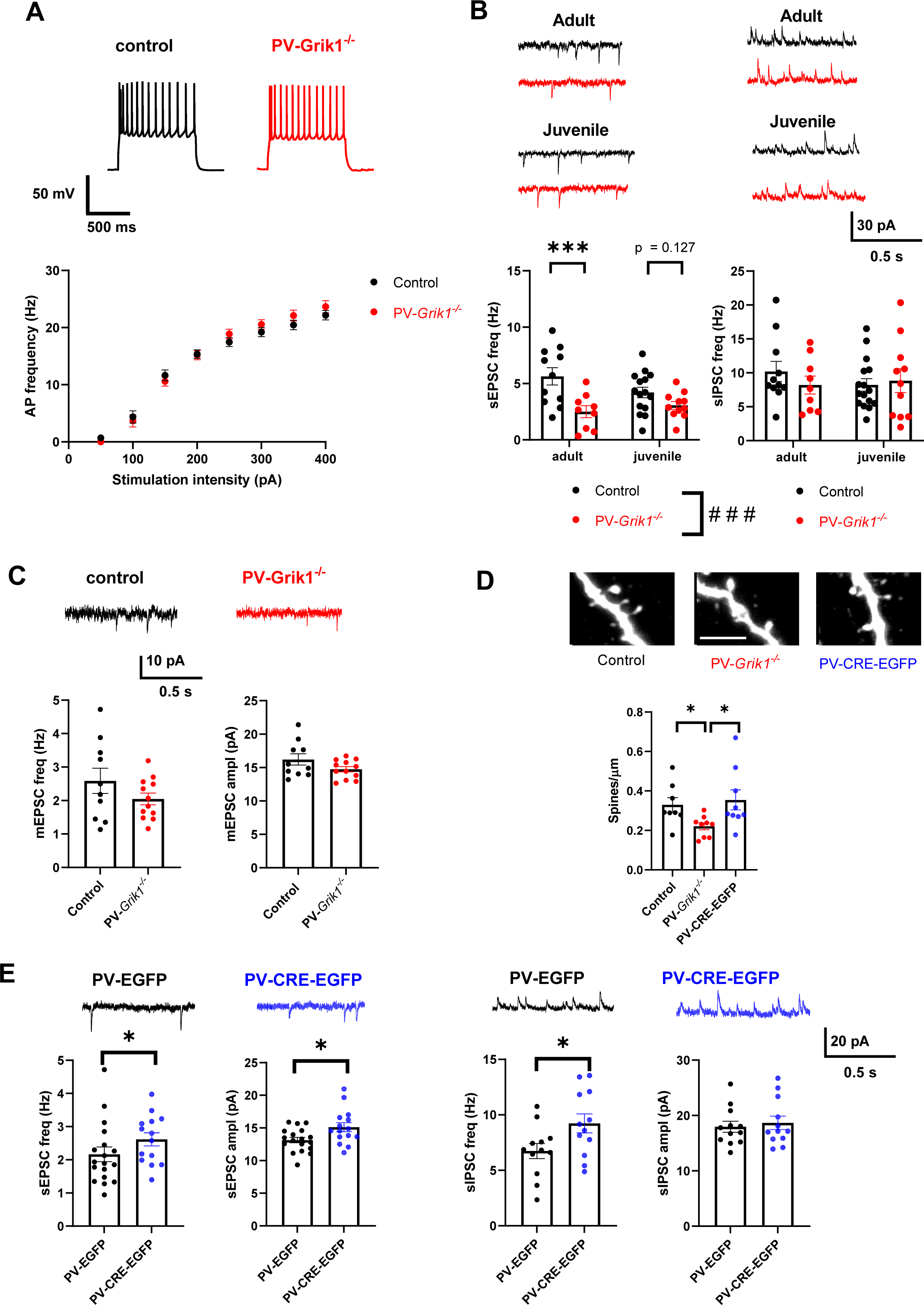
Synaptic inputs and excitability of BLA principal neurons (PNs) in mice lacking GluK1 in BLA PV interneurons. **A.** BLA neuron excitability. Example traces (top; control: black trace, *PV*-*Grik1^−/−^:* red trace), as well as quantification of AP firing frequency of BLA neurons in response to depolarizing current steps (bottom). No significant differences in AP frequencies were observed when comparing control and *PV*-*Grik1^−/−^* mice (F_(1, 131)_ = 0.9559, p = 0.33, mixed effects model). **B.** Example traces of sEPSCs and sIPSCs, recorded from BLA PN’s from both adult (n = 11 cells, 5 animals) and juvenile control (black traces) (n = 15 cells, 4 animals) and *PV*-*Grik1^−/−^* (red traces) mice (adult: n = 9 cells, 4 animals, juvenile: n = 18 cells, 4 animals) (top). Below, quantification of sEPSC frequency (left) and sIPSC frequency (right), ***p < 0.001; Holm-Sidak, ### p < 0.001; 2-way ANOVA. **C.** Example traces as well as quantification of mEPSC frequency and amplitude from adult control (black trace) (n = 10 cells, 4 animals) and *PV*-*Grik1^−/−^* (red trace) mice (n = 11 cells, 4 animals). **D.** Confocal images as well as quantification of spine density of adult BLA neurons of control (n = 8 slices, 3 animals), *PV*-*Grik1^−/−^* (n = 9 slices, 3 animals) and adult *PV-Grik1* KO (PV-CRE-EGFP, n = 9 slices, 3 animals) mice. Scale bar = 5 µm. * p < 0.05; one-way ANOVA. **E.** Consequences of adult PV IN GluK1 inactivation on glutamatergic and GABAergic drive onto BLA neurons. Example traces and averaged data for sEPSCs and sIPSCs, recorded from BLA PN’s from adult control (PV-EGFP, black traces) and *PV-Grik1* KO (PV-CRE-EGFP, blue traces) groups (top) (PV-EGFP, sEPSC: n = 18 cells, 3 animals, sIPSC: n = 12 cells, 3 animals, PV-CRE-EGFP sEPSC: n = 14 cells, 3 animals, sIPSC: n = 12 cells, 3 animals), * p < 0.05; unpaired t-test; # p < 0.05 Mann Whitney test.

Thus, despite the robust loss of PV IN excitability in the absence of GluK1 expression, the main effect observed in principal neurons in PV-*Grik1^−/−^* mice was a paradoxical decrease in glutamatergic input. Next, we investigated whether this intriguing reduction in glutamatergic drive onto BLA PNs in PV-*Grik1^−/−^* was driven by changes in network excitability or reflected altered glutamatergic connectivity. To this end, we recorded miniature excitatory postsynaptic currents (mEPSCs) from adult BLA PNs, while simultaneously filling the cells with biocytin for *post hoc* spine morphological analysis. mEPSC frequency and amplitude were not significantly different between the genotypes (frequency: F_(1, 17)_ = 1.315, p = 0.268, amplitude: F_(1, 17)_ = 2.275, p = 0.151), however, spine density was significantly lower (F_(1, 16)_ = 5.391, p = 0.035) in the *PV*-*Grik1^−/−^* as compared to controls (Figure 5C,D).

Maturation of PV IN function regulates closure of the critical periods of plasticity and might thereby affect development of the glutamatergic connectivity. To study whether the loss of glutamatergic connectivity to principal neurons in *PV*-*Grik1^−/−^* mice was of developmental origin, we used our adult knockout model. Interestingly, in *Grik1^tm^*^1c^ mice injected with the AAV-PV-CRE-EGFP viral vectors, both the frequency (p = 0.0399, Mann-Whitney) and amplitude (p = 0.0167, two tailed t-test) of sEPSCs in the BLA PNs was higher as compared to controls (Figure 5E). We also observed a slight increase in the frequency, but not the amplitude of sIPSCs (Figure 5E). Morphological analysis of the recorded neurons revealed no differences between the groups in spine density (F_(1, 16)_ = 0.153, p = 0.701, one-way ANOVA) (Figure 5D). Thus, acute and chronic (developmental) ablation of the GluK1 expression from PV INs has distinct effects on synaptic transmission and glutamatergic connectivity to BLA principal neurons.

Together, our data suggests that impaired PV IN function in the absence of GluK1 expression can be largely compensated for at the network level, to maintain the excitability and GABAergic drive of the BLA PN’s. In contrast, we detected developmentally originating persistent alterations in glutamatergic synaptic transmission and connectivity.

### Elimination of GluK1 in PV interneurons affects feedforward inhibition and synaptic plasticity in the BLA

PV INs are directly activated by cortical inputs, to mediate feedforward inhibition (FFI) in the BLA circuitry. In addition, PV IN excitability gates activation of the somatostatin interneurons, which disinhibits the circuitry mediating dendritic FFI in the BLA ^25,39^. Since PV IN excitability was significantly perturbed in *PV*-*Grik1^−/−^* mice, we sought to understand if FFI, directed at BLA PNs was also altered. To this end, we did whole-cell voltage clamp recordings from BLA PNs and recorded the ratio of evoked excitatory and disynaptic inhibitory postsynaptic currents (EPSC/dIPSC ratio), in response to stimulation of the cortical inputs to BLA. We found that FFI is significantly stronger in *PV*-*Grik1^−/−^* mice compared to controls (F_(1, 16)_ = 5.144, p = 0.039) (Figure 6A), suggesting that GluK1 ablation in the PV INs predominantly affected the BLA disinhibitory microcircuit.

**Figure 6.**
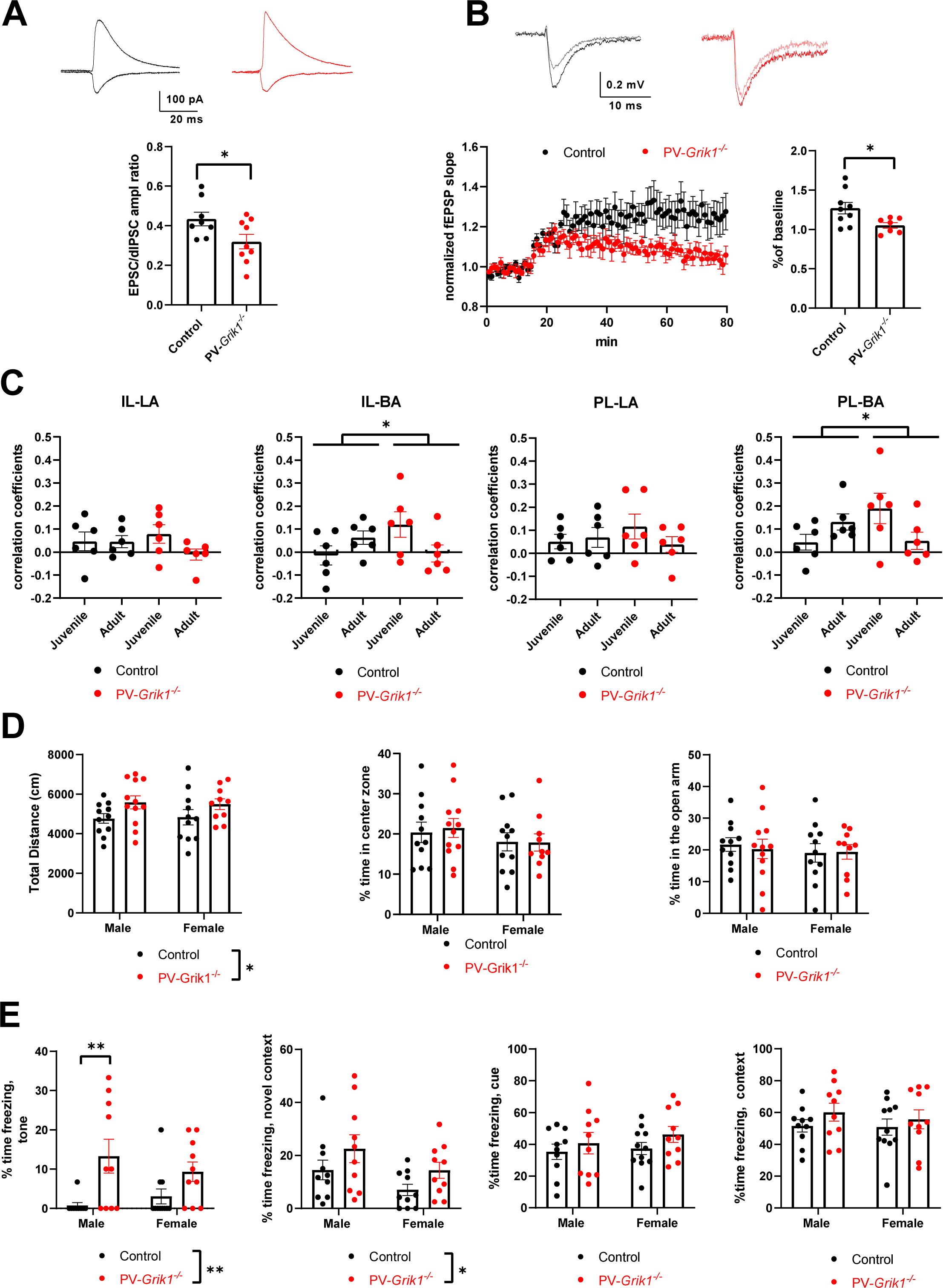
Absence of GluK1 in PV interneurons affects BLA microcircuit function, mPFC-BLA resting state functional connectivity and amygdala-dependent behaviors. **A.** FFI at cortical inputs to BLA PNs. On the top, example traces of EPSC and dIPSC, recorded from BLA principal neuron in control (black traces) and *PV*-*Grik1^−/−^* (red traces) mouse. Traces represent the average of 5 responses. On the bottom, quantification of EPSC/dIPSC ratio (control: n = 8 cells (4 animals), *PV*-*Grik1^−/−^*: n = 9 cells (6 animals); * p < 0.05, one-way ANOVA). **B.** LTP recordings. On top, example traces of fEPSPs during baseline (light gray and light red, for control and *PV*-*Grik1^−/−^*, respectively) and after 1 hour of TBS stimulation (black and dark red, for control and *PV*-*Grik1^−/−^*, respectively). Traces represent the average of 5 responses. TBS induced changes in synaptic strength were significantly different between genotypes (control: n = 9 slices (4 animals), *PV*-*Grik1^−/−^*: n = 7 slices (3 animals) * p < 0.05, one-way ANOVA). **C.** mPFC-amygdala resting state functional connectivity in juvenile (P21) and adult (P60) control and *PV*-*Grik1^−/−^* mice (control: n = 6, *PV*-*Grik1^−/−^*: n = 6, all males). * p < 0.05, significant genotype-age interaction, 2-way ANOVA. Values on the Y-axis indicate Pearson’s correlation coefficients. **D.** OF (left and middle) and EPM (right) behavioral tests in male and female *PV*-*Grik1^−/−^* mice with littermate controls. Total distance travelled in the OF was significantly different between the genotypes (* p<0.05; two-way ANOVA). No differences between the groups were detected in the % time spent in the center zone of the OF arena (F _(1, 40)_ = 0.04091, p = 0.8407, 2-way ANOVA) or in open arms in the EPM (F_(1, 40)_ = 1.072, p = 0.3067, two-way ANOVA). Control: male n = 11, female n = 11; *PV*-*Grik1^−/−^*: male n = 12, Female n = 10. **E.** *PV-Grik1^−/−^* mice exhibit a fear of novelty. Time spent freezing during a novel tone and in novel context was significantly different between the genotypes (* p < 0.05 ** p < 0.005, two-way ANOVA with Holm-Sidak). In fear conditioning, no differences were found in time spent freezing when presented with the conditioned cue (F_(1, 37)_ = 1.930 p = 0.1730) or when placed in a novel context (F_(1, 37)_ = 1.653 p = 0.2065). Control: male n=10, female n=11; *PV*-*Grik1^−/−^*: male n = 10, Female n = 10.

Since FFI is one of the key mechanisms controlling induction of synaptic plasticity in the BLA, we further wanted to study whether long-term potentiation (LTP) was disrupted in the *PV*-*Grik1^−/−^* mice. To this end, we performed extracellular field recordings from the lateral amygdala (LA) while stimulating the cortical afferents. Theta-burst stimulation (TBS) reliably induced a small but stabile synaptic potentiation in the control mice (Figure 6B). In contrast, in the *PV*-*Grik1^−/−^* mice, TBS resulted in short–term potentiation, after which the synaptic responses returned back to the basal level. The TBS induced changes in synaptic strength were significantly different between genotypes (F_(1, 15)_ = 5.975, p = 0.028) (Figure 6B).

Impaired LTP in the *PV*-*Grik1^−/−^* mice is consistent with lower glutamatergic drive and increased FFI directed at the LA principal neurons, both of which increase the threshold for activation of NMDA receptors.

### Absence of GluK1 in PV interneurons results in sex and age specific changes in mPFC-BLA functional resting state connectivity

Considering the significant perturbations in FFI and LTP in the cortico-amygdaloid inputs, we sought to understand how the absence GluK1 from PV INs affected development of functional connectivity between medial prefrontal cortex (mPFC) and amygdala. To this end we recorded resting state functional connectivity from P21 and P60, male and female, control and *PV*-*Grik1^−/−^* mice using functional ultrasound imaging (fUS). Interestingly, there were significant age- and sex-dependent differences between the genotypes between infralimbic (IL) and prelimbic (PL) areas of mPFC and basal amygdala (BA) (Figure 6C, Supplementary Figure 4). Thus, at P21 but not at P60, connectivity between both L-BA and IL-BA was higher in the *PV*-*Grik1^−/−^* male mice as compared to the controls. Furthermore, in contrast to controls, PL-BA and IL-BA resting state connectivity decreased during development in male *PV*-*Grik1^−/−^* mice (Figure 6C). Both effects were specific to males, as there were no significant differences in mPFC-BLA resting state functional connectivity between female control and *PV*-*Grik1^−/−^* mice at either developmental time point (Supplementary Figure 4). Additionally, we observed that the connectivity between mPFC and BLA was significantly higher in females as compared to males a both developmental time points studied (Supplementary Figure 4).

### Absence of GluK1 in PV interneurons results in hyperactivity and fear of novelty

Finally, we wanted to investigate the behavioral consequences ensuing from the loss of GluK1 in PV INs. We subjected littermate *Grik1^tm^*^1c^ and *PV*-*Grik1^−/−^* mice (heterozygous for *Cre*) to open-field (OF), elevated plus maze (EPM) and fear-conditioning (FC) behavioral assays. Interestingly, although both GluK1 and PV INs have been implicated in anxiety ^40^ the *PV*-*Grik1^−/−^* mice did not display a classical anxiety-like phenotype in the EPM or OF, as no significant differences between the genotypes were observed in the time spent in the open arm or center zone, respectively. However, the total distance travelled in the open field by *PV*-*Grik1^−/−^* mice was significantly higher than their littermate controls, suggesting a hyperactive phenotype (F_(1, 40)_ = 5.461, p = 0.0245, two-way ANOVA) (Figure 6D).

The FC paradigm revealed no impairments in contextual or cue-dependent fear learning. However, the *PV*-*Grik1^−/−^* mice displayed fear of novelty. % time spent freezing was not different between genotypes under basal conditions, but was significantly higher in *PV*-*Grik1^−/−^* mice as compared to the littermate controls in a novel context (F_(1, 36)_ = 4.324, p = 0.0448, 2-way ANOVA) and when presented with a novel tone (F_(1, 36)_ = 11.88, p = 0.0015, 2-way ANOVA) (Figure 6E). These behavioral results suggest crucial function of GluK1 in PV INs in brain circuits encoding novelty detection and fear, both of which are suggested to involve amygdala ^41^.

## Discussion

Kainate receptors are powerful regulators of GABAergic transmission in the adult brain^42,43^. While strongly implicated in development of glutamatergic synapses and excitatory circuitry^44^, less information exists on the role of KARs in the development and maturation of GABAergic interneurons^11–13^. Our present results indicate a pivotal role of GluK1 KARs in the specification of PV IN phenotype during development and in maintaining the PV IN specific features in the adult BLA.

Constituting the largest subpopulation of cortical interneurons (∼40%) ^45^, PV INs play crucial roles in encephalic functions. Interestingly, PV INs are also the most vulnerable population of interneurons^37,46,47^ and their dysfunction is hence implicated in a multitude of neuropsychiatric and neurological disorders^46,48,49^. PV INs exert their crucial functions and are distinctive by their unique properties such as high AP frequency, relatively low input resistance, and high-amplitude rapid after-hyperpolarization^15,36^. These characteristic electrophysiological attributes were lost when GluK1 expression was eliminated, which may point to the etiological mechanisms of neurodevelopmental disorders where both KARs and PV INs are implicated, including e.g. mood- and anxiety disorders and schizophrenia^25,26,50–52^.

### GluK1 expression regulates development and maintenance of the PV interneuron phenotype

While NMDA-R mediated signaling has been identified to be critical for maturation of PV INs^19,21,23,53–56^, no previous data on KARs in this context exists. Here, we confirm that GluK1 subunit containing KARs contribute to glutamatergic drive onto PV INs during postnatal development, before they are fully mature. In the absence of GluK1 signaling, PV INs failed to acquire their adult phenotype and were characterized by low PV expression levels and low excitability throughout development. Furthermore, GluK1 lacking PV INs failed to integrate into the surrounding glutamatergic network and received significantly less glutamatergic inputs (sEPSCs) in the adult stage as compared to controls.

Interestingly, this phenotype was not observed in the juveniles, where the sEPSC frequency in GluK1 lacking PV INs was slightly higher than in controls. We postulate that the elevated frequency of sEPSCs in juveniles reflects a developmentally restricted homeostatic compensatory mechanism to counter loss of GluK1 and consequent loss of PV IN excitability.

Remarkably, inactivation of GluK1 expression at adulthood was sufficient to induce significant PV IN dysfunction, involving loss of PV labeling, high frequency firing and glutamatergic input. Although these effects were not as robust as compared to *PV-Grik1*^−/−^ mice, lacking GluK1 expression in the PV INs throughout development, the finding suggests that GluK1 may regulate also the adult plasticity of PV INs ^57^. Dynamic changes in PV IN function in the adults has been implicated in cognitive processes such as memory encoding^57^ but also in the context of stress^57^ and drug actions^59^; however, the molecular mechanisms involved are not well understood.

The significant attenuation in PV IN excitability, coupled with the loss of electrophysiological signatures such as AP halfwidth and high amplitude AHP current point to aberrant expression or function of voltage gated Kv3 potassium channels in the absence of GluK1. Typically, PV IN maturation is accompanied by increased expression of Kv3 channels, which ensure fast action potential repolarization important for the high frequency firing^15,60,61^. The developmental reduction in AP-halfwidth was not observed in PV INs lacking GluK1. In addition, high amplitude afterhyperpolarization, characteristic for PV INs^15,36^ was also lost in the *PV-Grik1*^−/−^ mice. Activation of GluK1 KARs can regulate voltage-gated channels via G-protein coupled signaling, which is implicated in regulation of neurotransmitter release and neuronal excitability^1,3,11,62^. In particular, in immature hippocampal interneurons, G-protein coupled GluK1 KARs regulate the apamine-sensitive K+ currents responsible for the medium-duration afterhyperpolarization (mAHP), which directly influences the interneuron firing rate^11^. Our data suggest that the role of KARs in controlling voltage gated channels goes beyond these previously described acute modulatory effects. The present results are consistent with a scenario where GluK1 signaling, likely via a G-protein coupled mechanism, is necessary to maintain the repertoire of voltage gated channels responsible for the unique membrane properties and specification of the PV IN subtype.

Interestingly, both GluK1 and Kv3 channels are regulated by BDNF signaling via the TrkB receptors^59,63,64^, and the same signaling pathway is strongly implicated in both developmental maturation and adult plasticity of PV INs^17,59^.

### Absence of GluK1 in the PV interneurons causes broader amygdala dysfunction

Each PV IN makes multiple synaptic contacts to a vast number of principal neurons, so that even the perturbation of a single PV IN function could alter the surrounding network drastically^51^. Surprisingly, despite the robust loss of PV IN excitability, we detected no changes in the excitability or GABAergic input in the BLA principal neurons in the *PV-Grik1*^−/−^ mice. Apparently, the developmental aberrations in PV IN function may be largely countered by compensatory changes in the network to maintain amygdala excitability.

PV INs also form strong connections with other interneurons, forming disinhibitory microcircuits that modulate feedforward inhibition (FFI) of principal neurons in the BLA^37–39^. We have previously shown that the primary effect of GluK1 antagonism in the adult LA is an increase in FFI due to release of PV IN mediated inhibitory control of somatostatin interneurons^25^. Consistently, ablation of GluK1 in PV INs resulted in an increase in the strength of FFI at cortical inputs to LA principal neurons, confirming that PV IN GluK1 receptors are necessary for balancing FFI in the LA. Furthermore, consistent with the critical role of FFI in gating LTP induction^65,66^, LTP in the cortical inputs to BLA was significantly attenuated in PV-*Grik1^−/−^* mice. Thus, while the some of the expected effects of PV IN dysfunction on BLA circuit excitability appear to be developmentally compensated for, our data reveals an indispensable function for PV IN GluK1 receptors in controlling the strength of FFI and consequently, threshold for LTP induction in the BLA.

Intriguingly, we also observed that glutamatergic drive to BLA neurons was clearly diminished in juvenile and significantly reduced in adult PV-*Grik1^−/−^* mice. This phenotype was of developmental origin, as it was not seen in response to inactivation of PV IN GluK1 receptors in adults. Parallel changes in spine density further show that developmental loss, but not adult ablation of GluK1 expression in PV INs results in lower glutamatergic connectivity to the BLA principal neurons. PV INs play a significant role in the development and fine tuning of cortical circuits and are vital for the closure of the critical period^16^. Precocious maturation of PV INs during early life stress is thought to drive premature closure of the critical period plasticity and result in significant alterations in cortico-amygdaloid connectivity across development^65–68^. Since, in our model, PV INs don’t seem to reach full maturity in the absence of GluK1, this could perturb the closure of the critical period. Prolonged critical period plasticity involving excessive synaptic pruning could explain the reduced glutamatergic drive on to BLA neurons we observe in our knockout model.

Consistent with this scenario, we found that mPFC to BLA resting state connectivity is perturbed in PV-*Grik1^−/−^* mice. In control mice, there is a developmental increase in functional connectivity between mPFC and BLA. This is not only lost in the knockouts but is also reversed to a developmental decrease in functional connectivity and may be a consequence of excessive synaptic pruning. This supports the idea that developmental dysfunction of GluK1 lacking PV interneurons may indeed disturb the activity-dependent refinement and result in persistent changes in the cortico-amygdaloid connectivity.

### Behavioral outcomes of the PV interneuron specific GluK1 ablation

Behavioral testing revealed that the PV-*Grik1^−/−^* mice were hyperactive and had increased fear of novelty. In contrast, ablation of GluK1 selectively in PV INs was not sufficient to produce anxiety-like behavior, where these receptors have been previously implicated^7,25^. However, both anxiety and fear of novelty depend on amygdala circuitry^41^, highlighting a role for GluK1 in regulating amygdala dependent behaviors.

KARs have been previously shown to regulate hyperactivity and psychomotor agitation^52,67,68^. Our data extends these findings and suggests that this phenotype depends on KARs dysfunction in PV INs.

The existing research linking PV IN dysfunction to neuropsychiatric conditions, especially schizophrenia is extensive, comprising both of human and animal model studies^46,47,69–72^. A majority of these studies reveal not only a diminishment in PV IN density and PV expression in several brain structures^53,73–75^, but also a loss of glutamatergic synapses in PV INs^76^. A similar loss of PV expression is seen in our knockout model and our results fall in line with existing literature on glutamatergic input to PV INs in the context of these disorders. Given these intricate and indubitable links, the results of our study may therefore be useful in interpreting and dissecting the etiological mechanisms in neuropsychiatric disorders such as schizophrenia.

In summary, we have shown the significance of GluK1 KARs in the functional and neurochemical maturation of PV INs in the BLA. The PV IN dysfunction in the absence of GluK1 associate with perturbations in BLA network function and behavior. These results illuminates the role of GluK1 KARs and PV INs in shaping BLA networks and provide scope for designing effectual therapeutic methods for disorders that entail disrupted GluK1 and PV IN function.

## Acknowledgements

We thank Janne Sulku, Vootele Voikar, the personnel in the Mouse Behavioral Phenotyping Facility and in the Laboratory Animal Center for expert technical help. This study was financially supported by the Academy of Finland, Sigrid Juselius Foundation, Magnus Ehrnrooth Foundation and Finnish Cultural Foundation. Mouse Behavioral Phenotyping Facility is supported by Biocenter Finland.

## Author contributions

J.H. and R.S. carried out the vast majority of electrophysiological experiments and data analysis, and M.R. and G.Z. contributed to these data. J.H., R.S., S.O. and J.K.R. performed the immunostainings and imaging. R.S. and J.S. performed the viral injections. The manuscript was written by J.H., R.S. and S.E.L.

