## Supplementary Figure for "GluK1 kainate receptors are necessary for functional maturation of parvalbumin interneurons regulating amygdala circuit function"

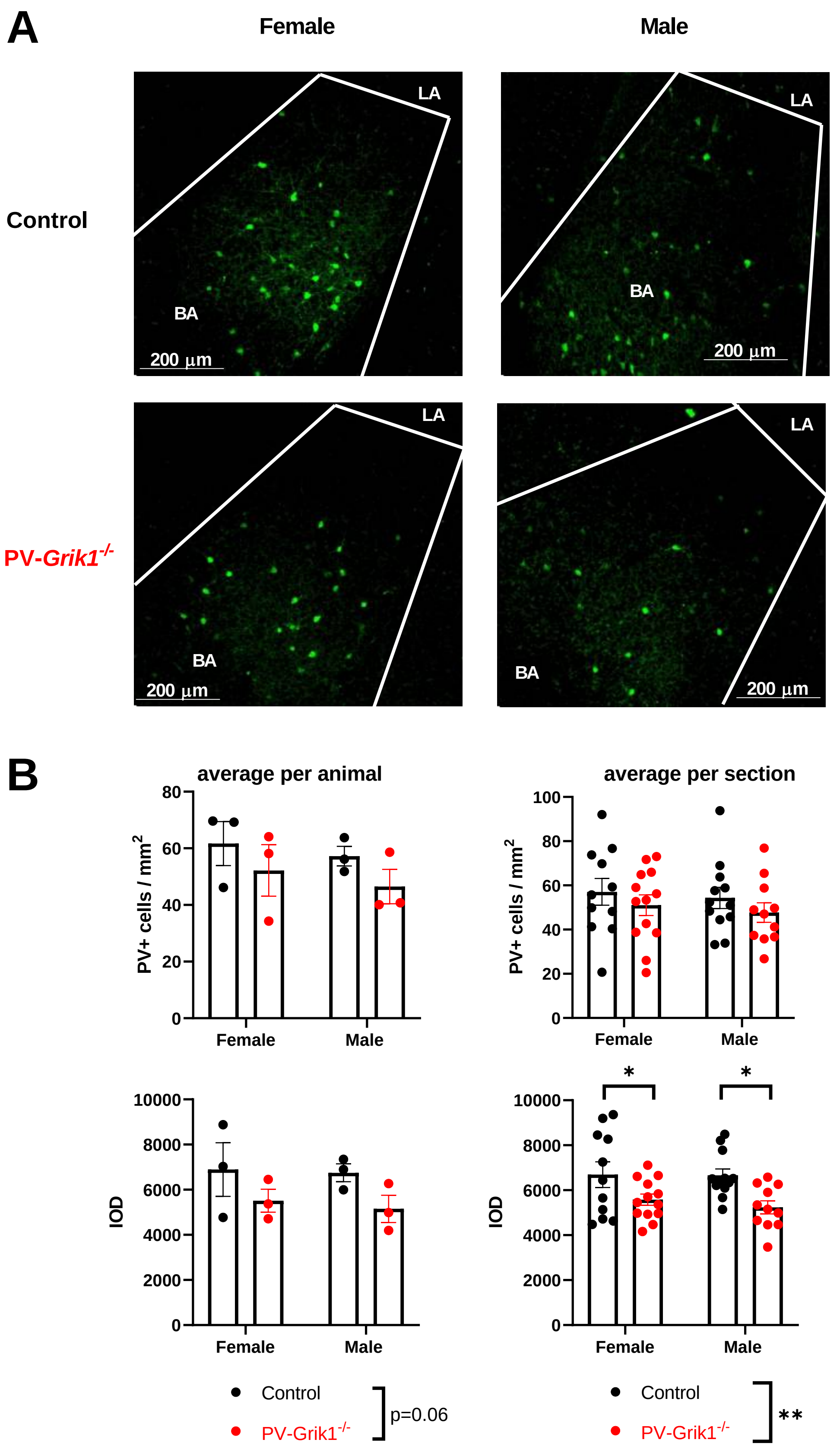


**Supplementary Figure 1**.

A. PV staining in female and male control and PV-Grik1-/- mice, that are heterozygous for *Cre*. Example pictures of PV (green) immunostaining in the BLA of adult female and male control (top row) and PV-Grik1-/- (bottom row) mice.

B. Quantification of the IOD and PV+ cell densities in both adult female and male BLA. The data consist of 3 animals per group and 3 - 5 slices per animal and are shown for both, data averaged per animal (left) (genotype effect, IOD: $F_{(1, 9)} = 4.542$, p=0.0619; 2-way ANOVA) and per section (genotype effect, IOD: $F_{(1, 43)} = 12.27$, p=0.0011;* $p<0.05$, 2-way ANOVA with Holm Sidak).

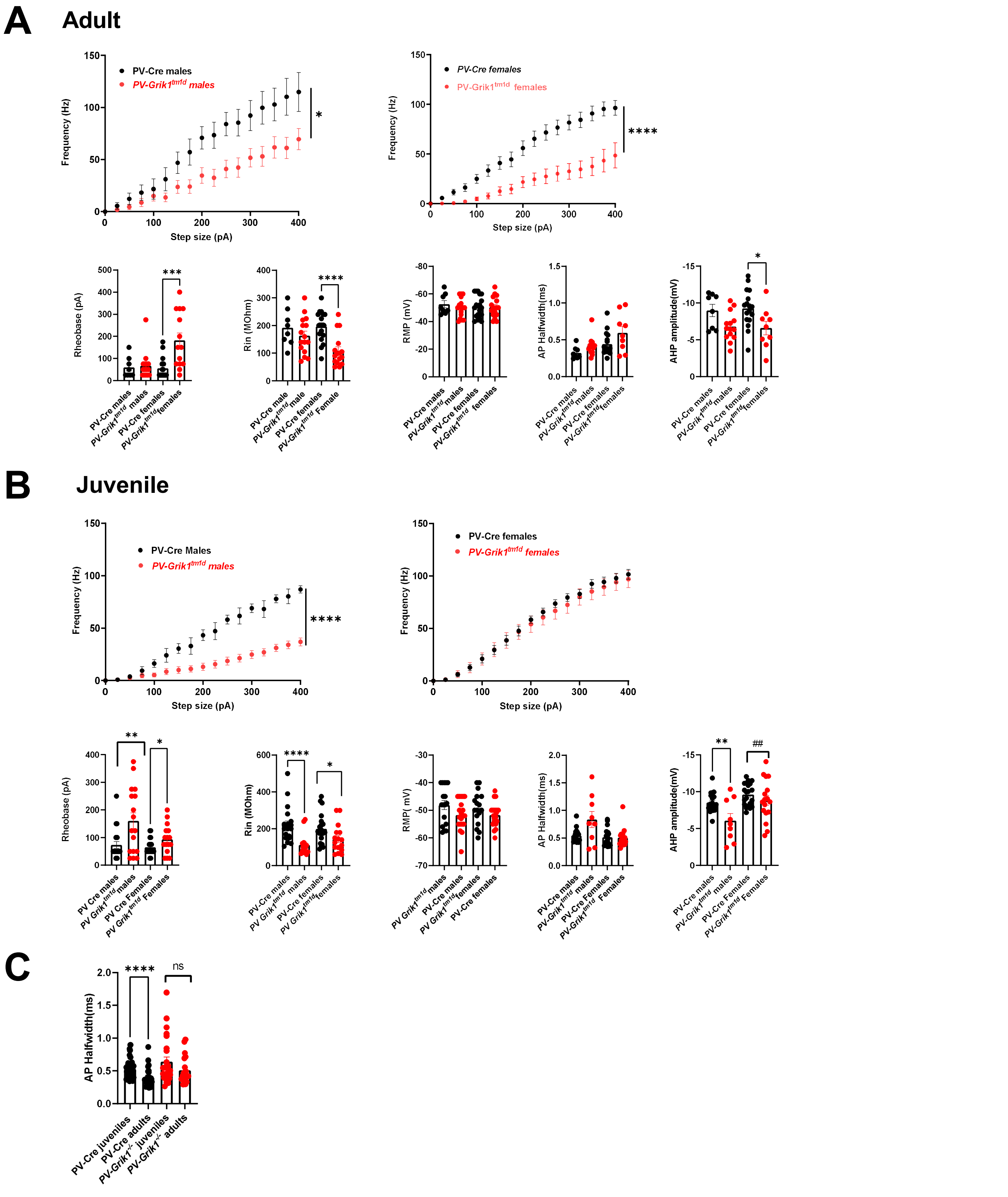


**Supplementary Figure 2. Absence of GluK1 results in low excitability of PV interneurons in both males and females across development.**

A. PV IN excitability data from adult, from the same recordings as in Figure 3A, analysed separately for males and females. Control males: n= 8 cells, 3 animals, females: n=18 cells, 5 animals; PV-Grik1-/- males: n=15 cells, 3 animals, females: n= 14 cells, 4 animals. * $p < 0.05$; ****$p < 0.0001$; mixed effects model. Quantification of rheobase, input resistance (Rin), resting membrane potential (RMP), AP halfwidth and AHP amplitude. * $p < 0.05$; ***$p < 0.001$; **** $p < 0.0001$; two-tailed t-test and Mann-Whitney tests .

B. Similar data from juveniles, from the same recordings as in Figure 3 G. Control males: n = 20 cells, 4 animals, females: n=21 cells, 4 animals; PV-Grik1-/- males: n= 17 cells, 4 animals, females: n=17 cells, 4 animals. *$p < 0.05$; **$p < 0.01$; **** $p < 0.0001$; ##$p < 0.01$; f-test for variances

C. Data comparing AP halfwidth between the age groups, for the same data as shown in A and B. **** $p < 0.0001$ Mann-Whitney test .

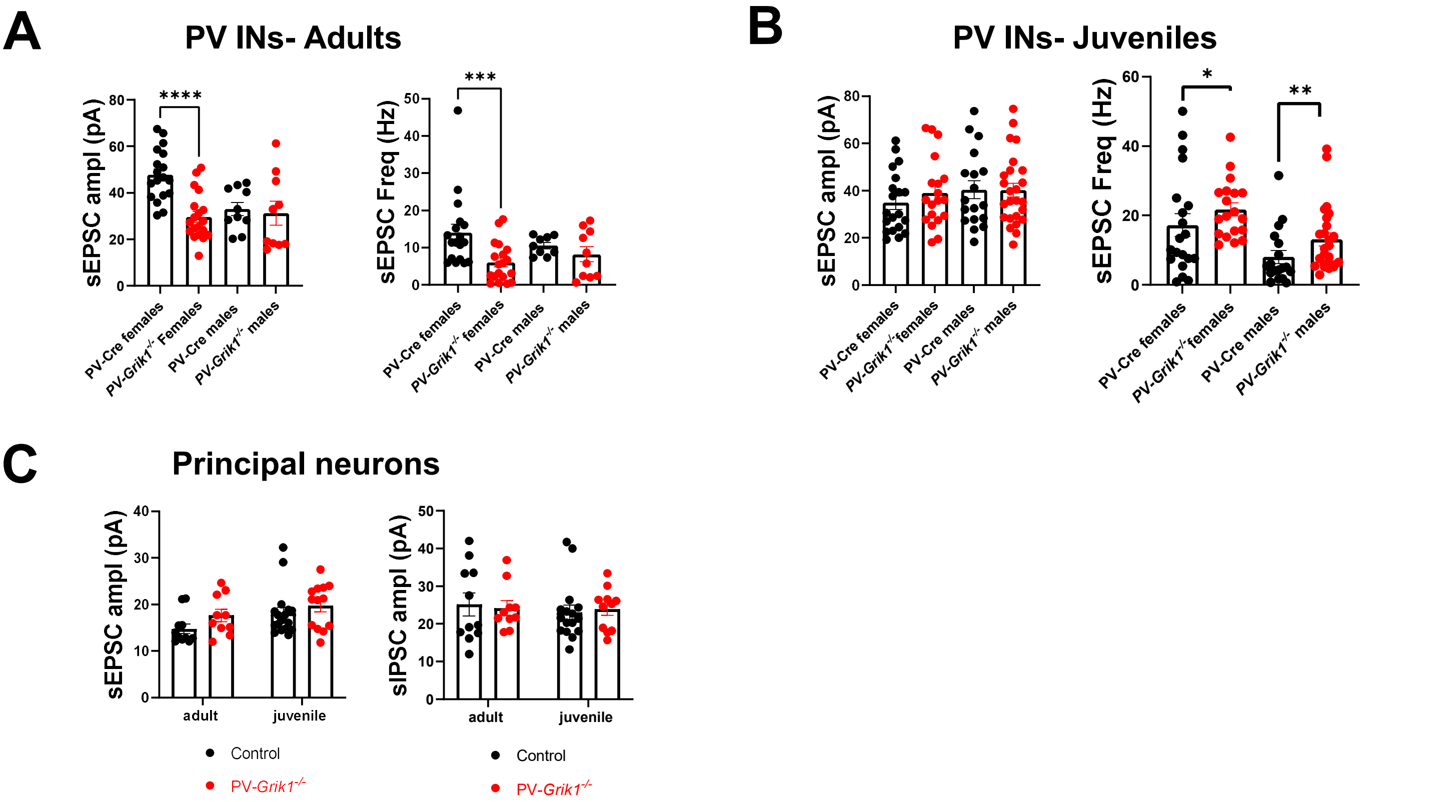


**Supplementary Figure 3.** Additional data related to analysis of spontaneous synaptic events in BLA PV INs (Figure 4) and principal cells (Figure 5B) in control and PV-Grik1-/- mice.

A. sEPSC frequency and amplitude data for the same adult PV IN recordings as shown in Figure 4A, analysed separately for males and females. Control females: n=18 cells 5 animals; males: n= 10 cells, 3 animals; PV-Grik1-/- females: n=19 cells, 4 animals; males: n=10 cells, 3 animals. ***$p < 0.001$; ****$p < 0.0001$; two-tailed t tests and Mann-Whitney test.

B. sEPSC frequency and amplitude data for the same juvenile PV IN recordings as shown in Figure 4B, analysed separately for males and females. Control males: n= 18 cells, 5 animals; females n= 20 cells, 4 animals; PV-Grik1-/- males: n=25 cells, 6 animals; females: n=19 cells, 4 animals.

C. sEPSC and sIPSC amplitude data for the same BLA principal neuron recordings that are shown in Figure 5B. Controls, Juvenile: n = 17 cells, 4 animals; adult: 11 cells, 5 animals; PV-Grik1-/- juvenile: n = 13 cells, 4 animals; adult: 10 cells, 5 animals.

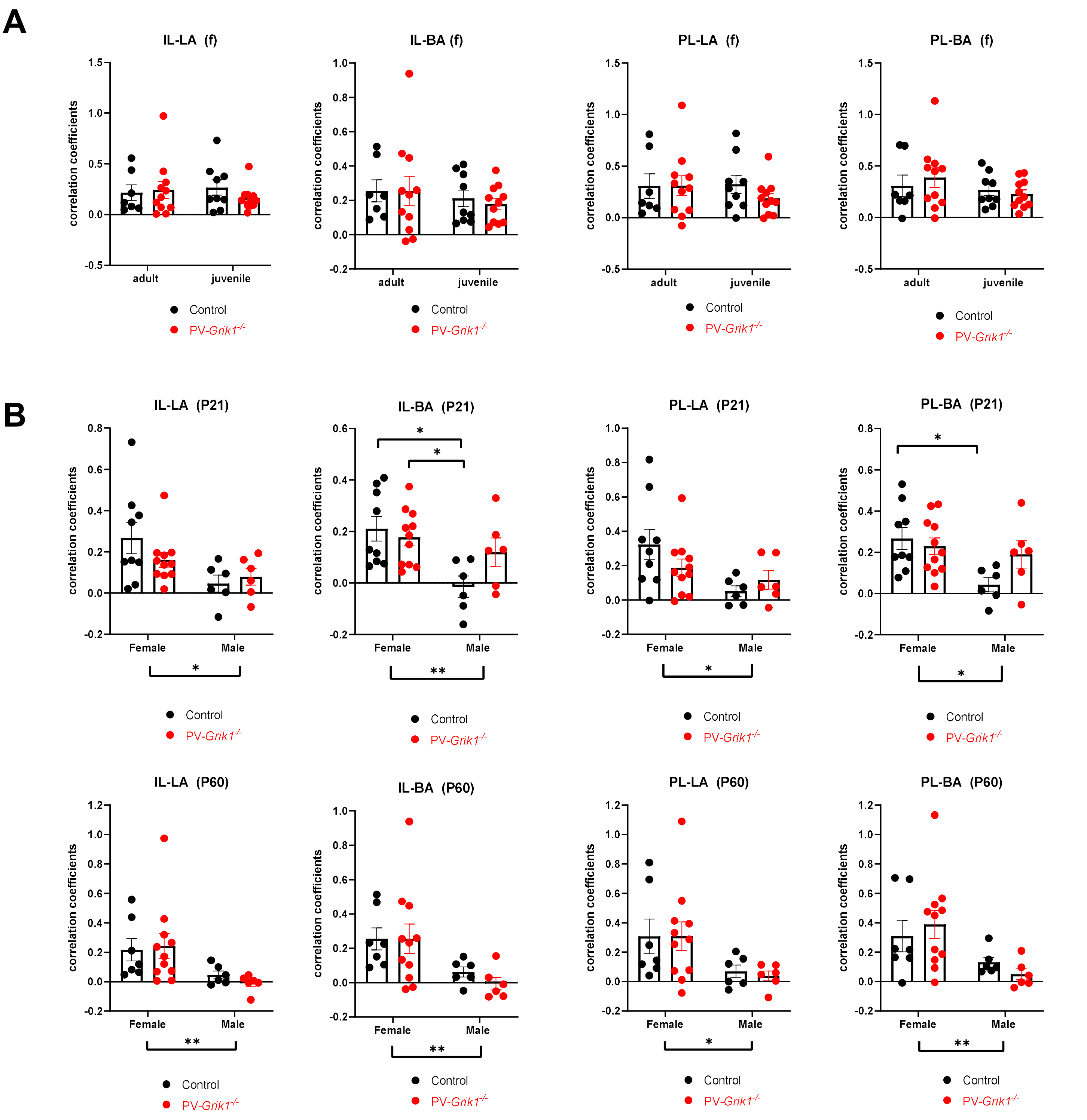


**Supplementary Figure 4.** Absence of GluK1 from PV INs results in age and sex specific changes in mPFC-BLA resting state functional connectivity.

**A.** fUS imaging data on resting state functional connectivity in adult (P60) and juvenile (P21) female control (n = 9) and PV-Grik1-/- (n = 11) mice. * $p < 0.05$, 2-way ANOVA. Similar data for males is shown in the Figure 6C.

**B.** Sex comparisons of resting state functional connectivity in juvenile (top row) and adult (bottom row) control and PV-Grik1-/- mice. The graphs are replotted from the same data as shown in Figure 6C and Supplementary Figure 4A. ** $p < 0.01$, *$p < 0.05$, 2-way ANOVA and Holm-Sidak. Values on the Y-axis indicate Pearson’s correlation coefficients.
